# *De Novo* Mutation in an Enhancer of *EBF3* in simplex autism

**DOI:** 10.1101/2020.08.28.270751

**Authors:** Evin M. Padhi, Tristan J. Hayeck, Brandon Mannion, Sumantra Chatterjee, Marta Byrska-Bishop, Rajeeva Musunuri, Giuseppe Narzisi, Avinash Abhyankar, Zhang Cheng, Riana D. Hunter, Jennifer Akiyama, Lauren E. Fries, Jeffrey Ng, Nick Stong, Andrew S. Allen, Diane E. Dickel, Raphael A. Bernier, David U. Gorkin, Len A. Pennacchio, Michael C. Zody, Tychele N. Turner

**Affiliations:** Department of Genetics, Washington University School of Medicine, St. Louis, MO 63110, USA; Department of Pathology and Laboratory Medicine, Perelman School of Medicine, University of Pennsylvania, Philadelphia, PA 19104, USA; Environmental Genomics and Systems Biology Division, Lawrence Berkeley National Laboratory, Berkeley, CA 94720, USA; Center for Human Genetics and Genomics, NYU School of Medicine, New York, NY 10016, USA; New York Genome Center, New York, NY 10013, USA; Center for Epigenomics. University of California San Diego School of Medicine, 9500 Gilman Drive, La Jolla, CA 92093, USA; Institute for Genomic Medicine, Columbia University, New York, NY 10027, USA; Center for Statistical Genetics and Genomics, Duke University, Durham, NC 27708, USA; Division of Integrative Genomics, Duke University, Durham, NC 27708, USA; Department of Biostatistics and Bioinformatics, Duke University, Durham, NC 27708, USA; U.S. Department of Energy Joint Genome Institute, Walnut Creek, CA 94598, USA; Department of Psychiatry and Behavioral Sciences, University of Washington, Seattle, WA, 98195 USA

## Abstract

Previous research in autism and other neurodevelopmental disorders (NDDs) has indicated an important contribution of *de novo* protein-coding variants within specific genes. The role of *de novo* noncoding variation has been observable as a general increase in genetic burden but has yet to be resolved to individual functional elements. In this study, we assessed whole-genome sequencing data in 2,671 families with autism, with a specific focus on *de novo* variation in enhancers with previously characterized *in vivo* activity. We identified three independent *de novo* mutations limited to individuals with autism in the enhancer hs737. These mutations result in similar phenotypic characteristics, affect enhancer activity *in vitro*, and preferentially occur in AAT motifs in the enhancer with predicted disruptions of transcription factor binding. We also find that hs737 is enriched for copy number variation in individuals with NDDs, is dosage sensitive in the human population, is brain-specific, and targets the NDD gene *EBF3* that is genome-wide significant for protein coding *de novo* variants, demonstrating the importance of understanding all forms of variation in the genome.

**One Sentence Summary:** Whole-genome sequencing in thousands of families reveals variants relevant to simplex autism in a brain enhancer of the well-established neurodevelopmental disorder gene *EBF3*.

## Introduction

Whole-genome sequencing (WGS) in large sample sizes is critical for understanding of the contribution of coding and noncoding variants in complex disease (*1*). In particular, this kind of data is essential for characterizing the contribution of variation in the enigmatic noncoding portions of the genome. When variants are found in these regions, different types of annotation based on items such as evolutionary conservation (*2*), constraint (*3*), and epigenetic marks (*4*) can help guide researchers to functionally important regions of noncoding DNA as a starting point for assessing variation within them. Genome-wide association studies (GWAS) have been critical for identifying noncoding variants that are common in the population and associated with human disease (*5*). However, WGS has now provided access to rare and *de novo* noncoding variants which are difficult to associate with phenotype without using aggregation methods (*6-9*); these methods have been shown to work for rare coding variation (*10-14*), but have been challenging to apply to non-coding regions because of the lack of clearly defined, discrete elements and sequence-based models of variant effect.

Autism is a complex neurodevelopmental disorder with a heritability of ∼80% (*15*). Large copy number variants (*16-19*) and protein-coding *de novo* variants, identified by the successful whole-exome sequencing strategy (*10-14, 20, 21*), contribute to ∼30% of cases with higher enrichment in females with autism and those with intellectual disability (*10*). Recently, we and other researchers have identified an overall enrichment of *de novo* (*6-9, 22*) or paternally inherited variants (*23*) within regulatory sequence in individuals with autism. However, these studies have mostly assessed this aggregation of genetic burden across a large panel of pooled regulatory elements. To begin to parse out the underlying biology of autism *de novo* variants in individual regulatory regions, we turn to the extensively studied VISTA enhancer dataset (*24, 25*). These enhancers were identified based on a variety of features (e.g. sequence conservation, epigenetic signatures, etc.) and have been tested in an embryonic mouse assay giving information on the spatial-temporal dynamics of the enhancer activity. Another recent study has also highlighted the utility of the VISTA dataset in identifying enhancers relevant in neurodevelopment (*9*). We adapted a model (*26*) for assessing excess *de* novo mutation load in protein-coding regions so that we could test for excess of variation in each of the VISTA enhancers with known ability to drive expression in the embryonic brain. Application of this test in 2,671 families with autism (n = 9,831 individuals) revealed VISTA enhancers with excess of *de novo* mutation in autism. We extensively tested the enhancer hs737 in follow-up genomic, epigenomic, clinical, *in silico, in vitro*, and *in vivo* analyses. While much work remains in understanding the role of noncoding variation, these analyses provide an initial framework for characterizing single enhancers in complex disease.

## Results

### de novo variant data in 2,671 families

The New York Genome Center has focused on whole-genome sequencing analyses to understand the role of all forms of variation in autism (*6-8, 22, 23, 27, 28*). We aggregated *de novo* variant data from a recent publication (*29*). The *de novo* variant data from that study (*29*) was downloaded through SFARI Base (See methods). In total, there were 292,407 *de novo* variants (DNVs) in 2,671 probands and 1,818 unaffected siblings.

### Statistical assessment of DNVs in VISTA elements

To assess for enrichment of DNVs, we modified the fitDNM (*26*) approach, a test of excess *de novo* load previously used across genes, to work on noncoding DNA sequences. In particular, we focused on 544 human elements (Supplemental table S1) that show enhancer activity in the brain using a lacZ transgenic assay at embryonic day 11.5 in mice (*25*). Of these, we tested 29 enhancers with at least one proband DNV (Supplemental table S2). There were 2 enhancers with 3 DNVs, 5 with 2 DNVs, and 22 with 1 DNV. Six enhancers exhibited excess of *de novo* variation (p < 0.05) (Table 1). One enhancer, with 3 DNVs, was hs737 (p = 1.12 × 10^−4^), which we had previously identified in the first 516 families as an enhancer with a gain-of-function DNV in an individual with autism (11257.p1) (*6*). The hs737 enhancer drives expression in the midbrain and hindbrain at embryonic day 11.5 (E11.5) (https://enhancer.lbl.gov/cgi-bin/imagedb3.pl?form=presentation&show=1&experiment_id=737&organism_id=1). The second enhancer, with 3 DNVs, was hs2333 (p = 3.97 × 10^−4^) and it drives expression in the forebrain and hindbrain at E11.5 (https://enhancer.lbl.gov/cgi-bin/imagedb3.pl?form=presentation&show=1&experiment_id=2333&organism_id=1). It has recently been shown to be an enhancer interacting with the promoter of *DYRK1A* in the human hippocampus (*30*); a gene known to be genome-wide significant for protein-coding variants in autism (*31, 32*).

**Table 1.**
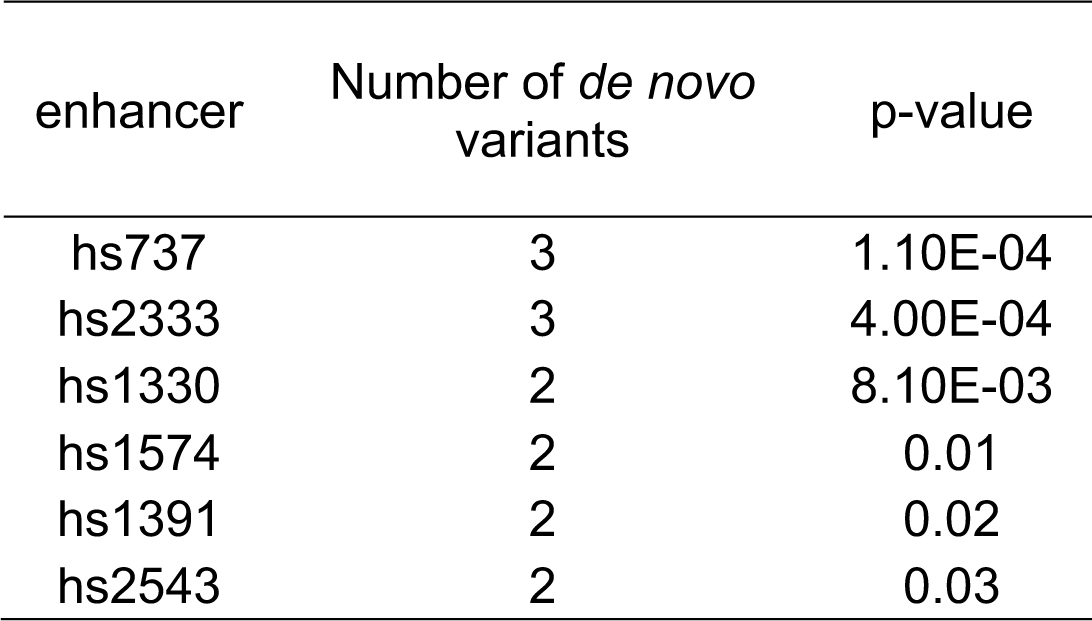
VISTA enhancers with an excess of *de novo* mutation based on fitDNM analysis.

| enhancer | Number of *de novo*<br>variants | p-value |
| --- | --- | --- |
| hs737 | 3 | 1.10E-04 |
| hs2333 | 3 | 4.00E-04 |
| hs1330 | 2 | 8.10E-03 |
| hs1574 | 2 | 0.01 |
| hs1391 | 2 | 0.02 |
| hs2543 | 2 | 0.03 |

### Dosage sensitivity of hs737 in the human genome

While *DYRK1A* is a plausible target of variants in hs2333, the putative target of hs737 was not immediately clear. Thus, we sought to better characterize hs737, and its potential gene targets. We hypothesize that if heterozygous point mutations in hs737 alter phenotype strongly, the enhancer would be dosage sensitive and show copy variation only in individuals with NDD. To test this, we first applied two approaches to assess copy number variation (CNV) in hs737. We assessed the morbidity map (*33, 34*) database containing 29,085 individuals with neurodevelopmental disorders (NDDs) and 19,584 controls. In particular, we looked at the window-analysis in Coe et al. 2014 (*33*) across the genome to identify the window containing hs737. This enhancer resided in a genomic window with an excess of deletions (case counts = 27, control counts = 0, p = 9.14 × 10^−7^) and duplications (case counts = 6, control counts = 0, p = 4.55 × 10^−2^) in individuals with NDDs. None of the 19,584 control individuals contained a CNV in this enhancer.

We also applied a tool to identify copy number (with paralog-specific sensitivity) in 1 kbp windows across the whole genome (*35*). We applied this to our WGS data and only identified one CNV in this enhancer. It was a deletion and occurred in proband 14091.p1 and upon further inspection was found to be a part of a larger known deletion (hg38: chr10:126450330-133655780 (*36*). To determine frequency of deletion/duplication in this enhancer in individuals without autism we also ran this copy number approach on the newly generated 3,202 individuals (high-coverage WGS) from the 1000 genomes project (http://ftp.1000genomes.ebi.ac.uk/vol1/ftp/data_collections/1000G_2504_high_coverage/). Combining our WGS parental data, 1000 genomes project data, and morbidity map there are no deletions or duplications in this enhancer in 28,128 non-NDD individuals (56,256 alleles) supporting this enhancer is sensitive to dosage in the human population. We also surveyed the gnomAD (*37*) database (v2.1) and observed no CNVs in the 10,847 individuals contained there (n = 21,694 alleles). To avoid possible double counting between gnomAD and other datasets, we do not present the aggregate data. Taken together, these results suggest that in addition to point mutations, structural variants involving hs737 may further contribute to NDDs.

### Clinical phenotypes of individuals with DNV in hs737

In order to understand more about the phenotypic consequences of variation in the enhancer, we reviewed clinical information for the each of the individuals with autism that had a DNV in hs737. As gene discovery in simplex autism, based on large CNVs and protein-coding DNVs, has been highest in females and in individuals who have intellectual disability (full scale IQ < 70) (*10*) the first two phenotypes that we assessed were sex and the full scale IQ scores. All three individuals were male and their full scale IQs were 103, 89, and 132, respectively (Figure 1A). This indicates that none of the individuals had intellectual disability. We also found that all three individuals had evidence of motor problems and/or hypotonia.

**Fig. 1.**
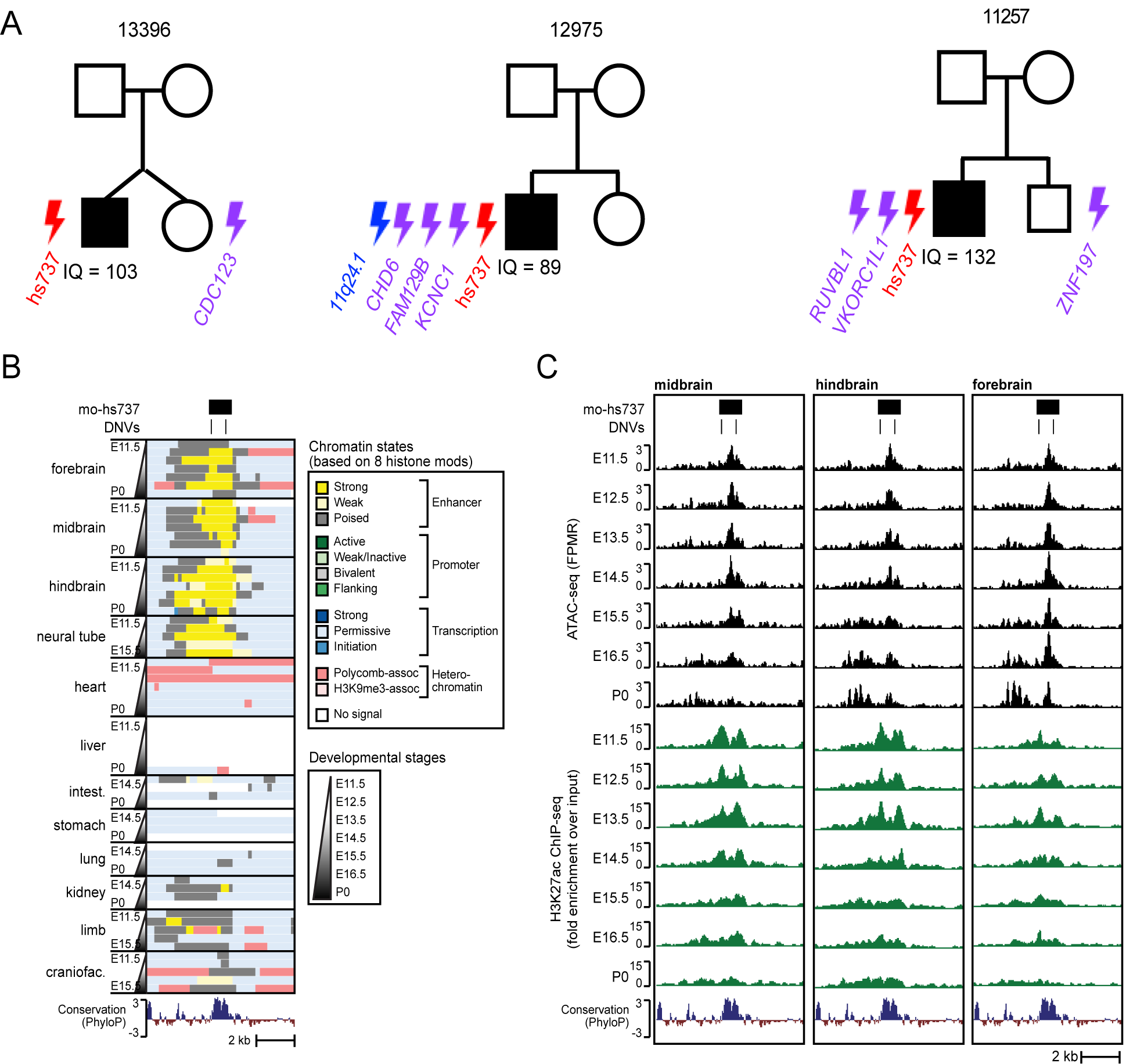
Individuals with *de novo* variation in hs737 and epigenetic characteristics of hs737. (A) Pedigrees of families with *de novo* variants in hs737. Lightning symbols indicate *de novo* variants with red = regulatory, purple = missense, and blue = deletion. Family identifiers are shown above the pedigree and the full scale IQ is shown below each proband. (B) Genome browser view (chr7:136,079,964-136,087,591; mm10) of chromatin states in mouse from (*40*) called by chromHMM (*41*) based on eight histone modifications: H3K4me1, H3K4me2, H3K4me3, H3K27ac, H3K27me3, H3K36me3, H3K9me3, H3K9ac. (C) Genome browser view (chr7:136,079,964-136,087,591; mm10) of ATAC-seq and H3K27ac ChIP-seq signal in midbrain, hindbrain, and forebrain at multiple developmental mouse stages from E11.5 to the day of birth (P0).

### Other genomic variation in individuals with DNV in hs737

In each individual with a DNV in hs737, we assessed the rest of the genome for other potentially relevant genomic variants that could be contributing to autism in each of the individuals (Supplemental table S3,S4). Individual 13396.p1 had no other large *de novo* CNVs or protein-coding *de novo* SNVs/indels. Individual 12975.p1 had three total *de novo* missense variants with one in each of the following genes *CHD6, FAM129B*, and *KCNC1* and also had a 1.6 Mbp deletion at 11q24.1 containing the following genes *BLID, BSX, C11orf63, CRTAM, SORL1*, and *UBASH3B*. Individual 11257.p1 had two *de novo* missense variants in each of the following genes *RUVBL1* and *VKORC1L1*. We scored each of the variants using a clinical scoring program (https://franklin.genoox.com/clinical-db/home) and all of the variants were classified as variants of uncertain significance.

### Epigenetic characteristics of hs737

Enhancers have well characterized epigenetic signatures that are predictive of their activity in specific biological contexts. Thus, to examine the activity of hs737 in its native genomic context, we took advantage of available epigenomic datasets from relevant human samples. We found that hs737 has several hallmarks of neuronal enhancer activity in humans including H3K27ac enrichment in fetal brain tissue (*38*), DNaseI hypersensitive sites (DHS) indicative of chromatin accessibility in CNS tissues (*6, 22*), and conserved transcription factor binding sites (TFBS) (*6*) (HMR Conserved Transcription Factor Binding Sites track in the UCSC Genome Browser (*39*)) (Supplemental Figure S1).

Hs737, and most other elements in the VISTA database, are assessed for reporter activity in mouse embryos at E11.5. To further examine the activity of this element during development, we analyzed the orthologous region in the mouse genome (mouse ortholog of hs737, or mo-hs737) using epigenomic data from a mouse embryonic developmental time series recently published (*40-42*). Consistent with the reporter expression pattern of hs737, we found that mo-hs737 has the chromatin signature of an active enhancer in midbrain and hindbrain at E11.5, but not in the non-neuronal tissues assayed (Figure 1B). Interestingly, mo-hs737 has the chromatin signature of a poised enhancer in forebrain at E11.5, but acquires the signature of an active enhancer at later developmental timepoints. This suggests that hs737 may have enhancer activity that extends to brain regions beyond the midbrain and hindbrain, and may explain why we previously saw gain-of-expression in the forebrain with one of the variants (*6*).

Strikingly, we found that characteristics of enhancer activity at mo-hs737 such as H3K27ac and chromatin accessibility (as measured by ATAC-seq) reach their height in brain tissue in mid to late gestation, and decline at birth (Figure 1C). This suggests that hs737 / mo-hs737 may exert its regulatory influence specifically during embryonic development, which could explain its involvement in NDDs, and may point to developmental stages and model systems that are most appropriate for future studies of this element.

### Gene target of hs737

To determine the potential gene target of the hs737 enhancer we first looked at all of the genes residing within the same topologically-associating domain (TAD) (hg38, chr10:128151746-130191746) (*43*). We focused on these genes since they are the most likely to be the targets of this enhancer. The genes included *C10orf143, CTAGE7P, EBF3, GLRX3, LINC01163, LOC728327, MGMT*, and *MIR4297;* the only gene in this region that is constrained in the human population with a gnomAD (*3*) o/e value for loss-of-function SNVs of 0.03 is *EBF3*. To examine physical interactions between hs737/mo-hs737 and genes in this region, we turned to high-resolution Hi-C data generated in a differentiation course from mouse embryonic stem cells (ESCs), to Neural Progenitor Cells (NPCs), and then to Cortical-like Neurons (CNs) (*44*). We found that the *Ebf3* promoter makes strong contacts across a large region encompassing mo-hs737, and that these interactions become stronger during differentiation to NPCs and CNs (Figure 2A). We used HiCCUPS (*45*) to call loops at each stage, and found that in CNs there is a ∼1.3 Mbp loop that brings mo-hs737 into close proximity with the *Ebf3* promoter (loop anchors chr7:136,050,000-136,075,000 and chr7:137,300,000-137,325,000; FDR=1.97 × 10^−18^). We did not observe loops between mo-hs737 and any other genes on the chromosome.

**Fig. 2.**
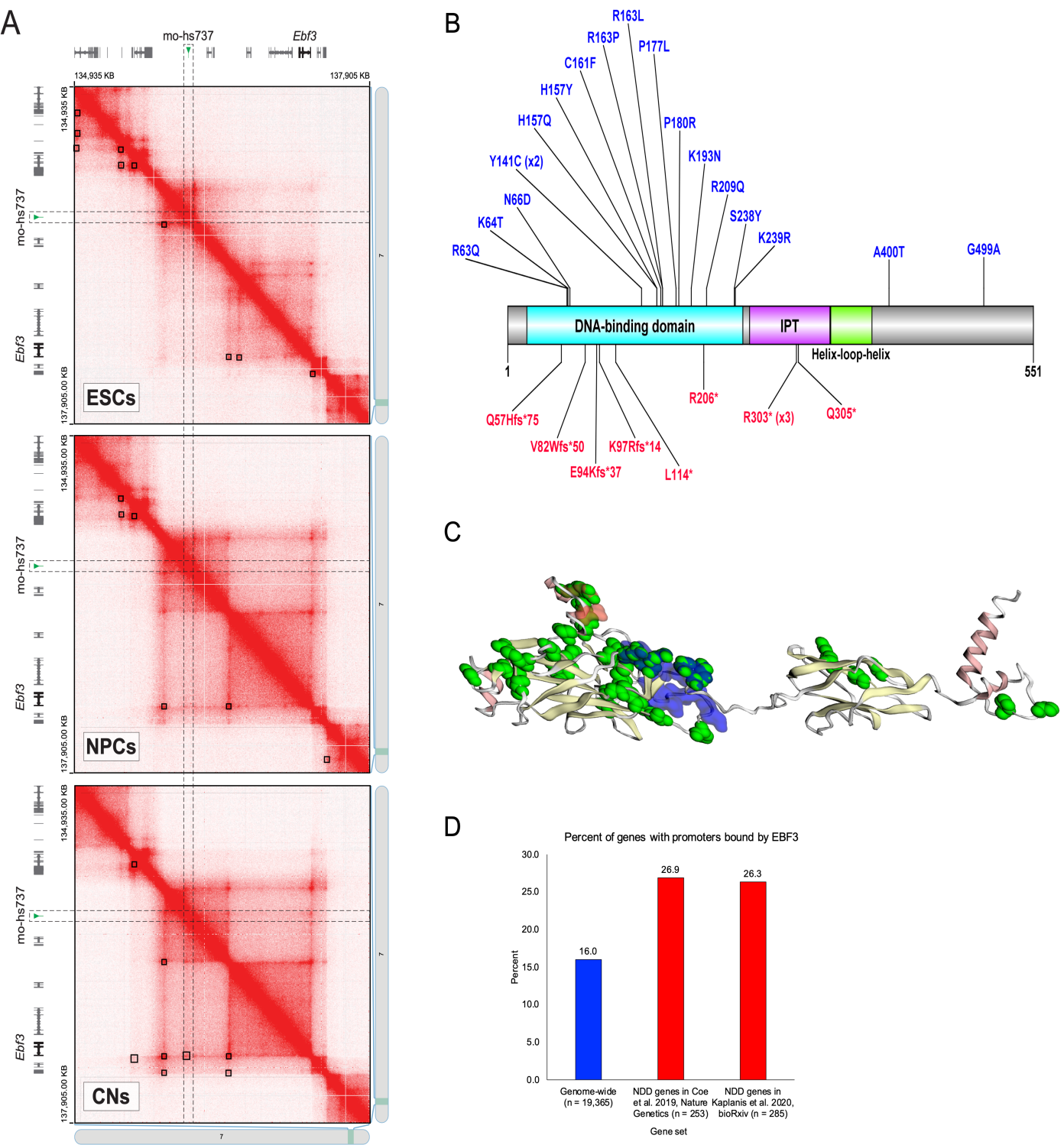
*EBF3* is the gene target of hs737. (A) Hi-C contact maps from Bonev et al. (*44*) visualized with Juicebox (*45*) at 5 kbp resolution. Heatmaps are symmetrical across the diagonal, except that HiCCUPS loop calls are shown as black boxes in the lower left half of each heatmap. The position of mo-hs737 is highlighted by dashed outlines. (B) EBF3 protein diagram (plotted using the DOG protein plotter (*66*)) with *de novo* variants identified in NDDs. Shown in blue are the missense variants and in red are the loss-of-function variants. (C) 3D model (plotted using the MuPIT program (*67, 68*)) of the EBF3 protein with *de novo* mutations identified in individuals with NDDs shown in green. (D) Genes with promoters bound by EBF3 based on ChIP sequencing in SK-N-SH cells. Enrichment is seen for the promoters of known NDD genesets.

Thus, narrowing in on *EBF3*, we searched recent literature on protein-coding DNVs and found *EBF3* is the only gene in this TAD that has known statistical enrichment for protein-coding DNVs in NDDs (*32, 46*). Combining data from these two previous studies (Figure 2B, Figure 2C, Supplemental table S5) we also saw significance (chimpanzee-human (*46*) missense p = 8.12 × 10^−35^, chimpanzee-human loss-of-function p = 2.26 × 10^−13^, denovolyzeR (*47*) missense p = 4.79 10^−13^, denovolyzeR loss-of-function p = 7.97 × 10^−22^) for protein-coding DNVs using two different statistical tests. This is also the gene for Hypotonia, Ataxia, and Delayed Development Syndrome (HADDS (*48*)) and has been characterized in detail in a set of ten patients (*49*).

### EBF3 regulates many NDD-significant genes

A recent publication (*50*) performed ChiP-sequencing on the EBF3 transcription factor in the SK-N-SH neuronal cell line. We mapped the peaks from this study to promoters in the human genome. In total, the promoters of 3,100 genes (16%) were bound by EBF3. We then focused in on genes with statistical enrichment for protein-coding DNVs in NDDs and that were bound by EBF3 (Supplemental table S6). Of the 253 significant NDD genes in Coe et al. 2019 (*46*), 26.9% of them were bound by EBF3 (p = 8.95 × 10^−6^, OR = 1.93) (Figure 2D). Of the 285 significant genes in Kaplanis et al. 2020 (*32*), 26.3% of them were bound by EBF3 (p = 7.43 × 10^−6^, OR = 1.87) (Figure 2D). Many of these bound NDD genes (Supplemental table S6) are involved in chromatin regulation (Chromatin Binding Gene Ontology p = 3.2 × 10^−7^) and/or transcription factor activity (DNA Binding Gene Ontology p = 6.4 × 10^−11^) (e.g., *CHD8, CHD2, ARID1B*) indicating that EBF3 may be a master-regulator of many NDD genes. Chromatin binding genes account for a sizeable fraction of DNM attributable cases of autism (*10, 12, 51*), suggesting that EBF3 disruption could result in a milder phenotype in the spectrum, as we observe in these cases.

### Mutagenesis of hs737 severely affects enhancer activity

To understand more about the hs737 enhancer and its resiliency to mutation, we designed two percent and five percent mutagenized alleles of the enhancer. We injected these into mice using a gold-standard lacZ transgenic assay (*52*). In this assay, we are able to determine the spatial-temporal dynamics of the enhancer. We found that these mutagenesis alleles sometimes inactivated the enhancer completely (Supplemental Figure S2), inactivated it partially (Supplemental Figure S2), or caused gain-of-function with new expression in other parts of the brain (Supplemental Figure S2). This indicates that many nucleotides of this enhancer are likely essential for appropriate enhancer activity.

### In vitro assessment of hs737 de novo mutations

In order to quantitatively assay the effects of hs737 *de novo* mutations identified in individuals with autism, we transfected the neuronal cell lines Neuro2a with a reporter construct that had the variant allele of the enhancer cloned upstream of a luciferase gene and a promoter (*53*). This design allowed for the assaying of quantitative effects of variation in a cell-type dependent manner. First, we examined the expression readout of the construct containing the known RET+3 enhancer variant (rs2435357) as a positive control (*53*) and found as expected that it had strong enhancer activity. In this cell line, the variants identified in individuals with autism displayed significant reduction in expression when compared to the reference allele and the promoter-only construct (Figure 3A, Figure 3B). This demonstrates that these variants within this enhancer can have quantitative effects.

**Fig. 3.**
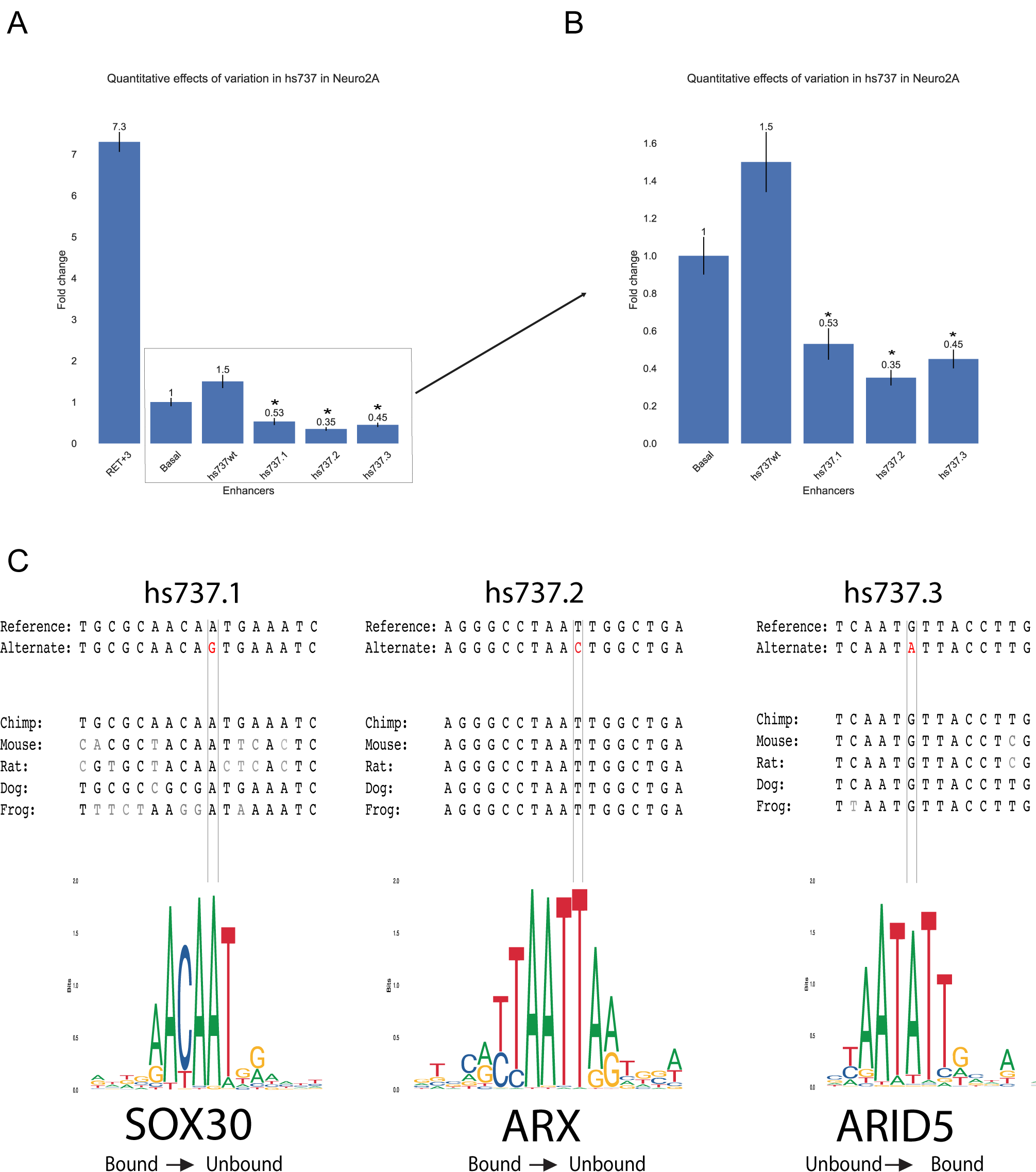
Effect of *de novo* variants, identified in individuals with autism, in hs737. (A) Results of luciferase assays in neuroblastoma (Neuro2a) cell lines with rs2435357 (RET+3) as a positive control for enhancer activity, promoter only (Basal), the wild type sequence of hs737 (hs737wt), and each of the three *de novo* mutations identified in individuals with autism. (B) Zoom in of A. (C) Sequence analysis of each of the three hs737 *de novo* mutations, identified in individuals with autism, including transcription factor binding site analysis results.

### In silico transcription factor binding assessment

To better understand functional consequences of the hs737 variants we observed, we analyzed the reference and variant enhancer sequences using QBiC (*54, 55*) (http://qbic.gcb.duke.edu), a program designed to predict the impact of noncoding mutations on transcription factor binding. We found that the *de novo* mutations, identified in individuals with autism, were each predicted to impact transcription factor binding (Figure 3C). Here we report the most significant hits, realizing that it could be any of the transcription factors in the family causing the functional impact on expression dynamics. All three variants are predicted to impact transcription factor binding via a transition mutation at a highly preferential base, that is also highly conserved (to frog). Each mutation occurs at a location within the position weight matrix that is predicted to completely change the binding status of the transcription factors. The first two variants are predicted to each respectively cause SOX30 and ARX to go from the bound state to unbound (SOX30 p = 2.48 × 10^−189^, z-score = −29.35; ARX p < 2.2 × 10^−16^, z-score = −40.27) while the third variant is predicted to create a new binding site for ARID5 (p < 2.2 × 10^−16^, z-score = 31.87). Mouse RNAseq at day E11.5 provides further support for these transcription factors being impactful as ARX and 12 members of the SOX family are in the 90^th^ percentile and the ARID family members are in the 80^th^ percentile of all genes that are expressed in the brain (Supplemental table S7).

## Discussion

Ten years ago, the Simons Simplex Collection began (*56*) and started its efforts to understand the role of genomic variation in simplex autism. It was hypothesized that there is a contribution from *de novo* variation in these simplex autism families (*56*). Over the ten years, microarray (*16, 18, 57*), WES (*10-12, 14*), and now WGS data (*6-8, 22, 23*) have been generated to fuel this discovery. The first fruits of these efforts were large copy number variants and protein-coding *de novo* variants. They have turned out to be critical for explaining ∼30% of individuals with autism (*10*). Intriguingly, these variants have been found to be enriched more in females and/or individuals who also had intellectual disability (*10*). Recent ongoing efforts looking at common variation are providing insights into other aspects of the genetics of autism and explain ∼50% of autism risk (*58*). In our study, we focused on the elusive *de novo* noncoding variation for which we and others have seen aggregate evidence for enrichment in promoters and enhancers (*6-8*). We also realize that we are under-powered in this arena until more genomes are generated in families with autism (*59*). For this reason, our approach focused only on the areas of the genome (enhancers) that are functionally validated to drive expression in the embryonic brain (in mice) and are also conserved in the human genome (i.e., VISTA enhancers) (*25*).

In our study, we identify and characterize the hs737 enhancer which has an enrichment of *de novo* mutations in autism. We find that beyond *de novo* single-nucleotide variants, copy number variants are also enriched in individuals with NDD. No control individuals have been identified that contain a deletion or duplication of this enhancer indicating that is dosage sensitive in the human population. These lines of genetic evidence support the finding that this enhancer has an important role in the human genome. Previous work in understanding the genetic contributions of *de novo* protein-coding variants have shown that there is an enrichment of individuals that are female and/or have intellectual disability (*10*) and that when examining individuals with mutations in the same gene shared phenotypes exist (*60, 61*). In our study, the individuals with noncoding DNVs in hs737 had shared phenotypes including being male, no intellectual disability, and all had hypotonia or motor delay. This was interesting since these phenotypes are more common than those seen in individuals with protein-coding DNVs and is likely because the noncoding variants have a smaller effect than protein-coding DNVs.

A major hurdle in the study of enhancers is determining which gene they regulate. This is especially relevant when enhancers are very distal from the promoter they target. Since hs737 resides in a large noncoding region, we utilized an innovative approach of combining constraint information for genes within the TAD, enrichment of protein-coding DNVs in genes in the TAD, and chromatin contact data. This data led to our identification of *EBF3* as the gene targeted by hs737 with an interaction across a distance of ∼1.4 Mbp. *EBF3* is a well-established NDD gene with genome-wide significant enrichment of protein-coding *de novo* variants and an established syndrome called HADDS that shares the phenotype of hypotonia with the individuals with autism in our study that had hs737 noncoding *de novo* variants. HADDS is a severe phenotype and affects many parts of the body. This is likely because EBF3 is a transcription factor that is ubiquitously expressed in humans (*62*). It affects many genes in the genome and in particular is enriched for regulating other NDD genes involved in further regulation. We speculate that since hs737 is a brain-specific enhancer that is why we see a less severe phenotype in the individuals with autism in our study than in HADDS. Each of the variants that we identified cause the enhancer to function differently than the wild type version of the sequence and all effect an AAT motif that is predicted to either disrupt or add a transcription factor binding site.

It is important to note that the road to further noncoding enhancer discovery in NDDs including autism will be difficult, requiring many genomes. Other studies have emphasized the overall enrichment of variation in noncoding elements (*6-9, 22, 23, 59*) with testing of specific variants in a small number of functional assays (*6, 8, 9*). However, each study has been limited and unable to reach genome-wide significance for all possible relevant elements in the genome due to the limited number of genomes that are available and the definition of the search space. Related to this issue, a gold-standard for multi-test correction p-values for noncoding elements is critical for future noncoding discovery much like the accepted standards for genome-wide association studies and analyses of whole-exome sequencing data. Next, we highlight the many resources that made a study like ours possible. If the decision had been made ten years ago to start with WGS in simplex autism, the finding of hs737 would not have been possible. In our study, we benefited from WGS efforts in autism led by the NYGC, the VISTA enhancer database carefully generated and curated at the Lawrence Berkeley National Laboratory, previous and ongoing efforts to identify all genes with protein-coding enrichment in NDDs using WES, constraint data in the human population (i.e., gnomAD (*3*)), and extensive characterization of the regulatory genome by ENCODE. Finally, we reflect on the phenotypes seen in individuals with hs737 *de novo* noncoding variants. All were male without intellectual disability and this presentation is the most abundant in simplex autism. Since, hs737 regulates EBF3 but has a weaker phenotype than in individuals with EBF3 coding mutations this potentially opens the door for brain-specific enhancer mutations in genes that are essential and therefore we never observe protein-coding mutations that are viable.

## Materials and Methods

### de novo variation in 2671 autism families

We downloaded *de novo* variant data from Wilfert et al. (*29*) through SFARI Base (accession: SFARI_SSC_WGS_2a, https://base.sfari.org/).

### Statistical assessment of *de novo* variants

A list of VISTA enhancers driving expression of their target genes in the brain were downloaded from the VISTA enhancer browser (*25*). *de novo* variants were annotated to each enhancer using bedtools (*63*). We modified the fitDNM statistical approach (*26*), a method to assess the excess mutational load of *de novo* variants using variant specific mutation rates calculated based on local sequence context, now applied to noncoding variants in the VISTA brain enhancers.

### Copy number assessment of hs737

To test copy number variant enrichment in morbidity map (*33*), we downloaded Supplementary Dataset 1 from Coe et al. 2014 (*33*) and identified the window in the genome containing hs737. We report in this paper the case counts, control counts, and p-values for deletions and duplications in this window.

The QuicK-mer2 (*35*) (https://github.com/KiddLab/QuicK-mer2) workflow was run on WGS data to generate copy number estimates, in 1 kbp windows across the genome, in each individual. Briefly, this method utilizes a kmer-based approach to perform copy number estimation. After running QuicK-mer2, we utilized the bedtools (*63*) map function to calculate the average copy number across the copy number windows covering hs737 (b38, chr10:128568604-128569741). If the copy number was less than 1.3, we called it as a deletion and if it was greater than 2.7, we called it as a duplication. QuicK-mer2 was run on the autism families in this study and also the high-coverage 1000 genomes project data available as described at http://ftp.1000genomes.ebi.ac.uk/vol1/ftp/data_collections/1000G_2504_high_coverage/.

To assess structural variation in gnomAD v2.1 (*37*), we queried for our enhancer region on hg19 (10-130366868-130368005) and also available at this link https://gnomad.broadinstitute.org/region/10-130366868-130368005?dataset=gnomad_sv_r2_1. There were “No variants found” in this region.

### Statistical testing of protein-coding DNVs in EBF3

*de novo* variant data was collected from two recent papers on NDDs (*32, 46*). After overlapping samples between the two studies were removed there were a total of 37,692 sequenced parent-child trios. To test for enrichment of protein-coding DNVs that were loss-of-function or missense, we applied the chimpanzee-human (*46*) and denovolyzeR (*47*) models as previously described (*46, 64*).

### Mouse ENCODE chromatin state and interaction tracks

Chromatin state and interaction data from mouse developmental timepoints were assessed in the ENCODE Regulation, ENC+EPD Enhc-Gene, ENCODE cCREs, and EPDnew Promoters tracks in the mm10 genome browser at UCSC (*39-42*).

### ChIP sequencing assessment

We downloaded ChIP sequencing data from Harms et al. 2017 (*50*) at the following GEO link https://www.ncbi.nlm.nih.gov/geo/query/acc.cgi?acc=GSE90682. To identify the promoters locations in he human genome, we looked at sequence 5 kbp upstream of the transcription start site using the Table Browser feature of the UCSC Genome Browser (*39*). We then used bedtools (*63*) intersect to identify which ChIP peaks overlapped with promoters in the human genome. To determine which NDD genes were bound at their promoter, we pulled the genome-wide significant gene lists from Coe et al. 2019 (*46*) and Kaplanis et al. 2020 (*32*) and compared to our EBF3 bound promoter list. Gene Ontology enrichment was performed using the Database for Annotation, Visualization and Integrated Discovery tool version 6.8 (https://david.ncifcrf.gov/) (*65*).

### RNA sequencing at mouse embryonic day 11.5

Mouse E11.5 forebrain RNAseq data was downloaded from https://www.encodeproject.org/files/ENCFF465SNB/@@download/ENCFF465SNB.tsv, mouse E11.5 midbrain RNAseq data was downloaded from https://www.encodeproject.org/files/ENCFF359ZOA/@@download/ENCFF359ZOA.tsv, and mouse E11.5 hindbrain data was downloaded from https://www.encodeproject.org/files/ENCFF750FTK/@@download/ENCFF750FTK.tsv. For each file, we retained all Ensembl gene identifiers and annotated them to HGNC approved identifiers using biomart (https://m.ensembl.org/info/data/biomart/index.html). We used an expression cutoff of > 2 to call a gene as expressed and < 2 as not expressed in each region of the brain.

### Random mutagenesis at E11.5 in mice

Random mutagenesis alleles were generated with either 2% or 5% of bases mutated in the enhancer. Each of these mutagenized enhancer sequences were assessed using a previously described transgenic lacZ reporter assay (*25*). Embryos with tandem integrations of the enhancer construct were assessed for expression differences from the baseline pattern seen in the wild type enhancer.

### Cell lines

Neuro2a (ATCC CCL-131) were purchased from ATCC and grown under standard conditions (DMEM + 10% FBS and 1% Penicillin Streptomycin).

### Luciferase assays

500 ng of firefly luciferase vector (pGL 4.23, Promega Corporation) containing the enhancer sequence cloned upstream of *luc2* and 9 ng of Renilla luciferase vector (transfection control) were transiently transfected into the Neuro2A cell line (1 × 10^5^ cells/well) using 3ul FuGene HD transfection reagent in 100ul of OPTI-MEM medium. Neuro2A Cells were incubated for 48 hours and luminescence measured using a Dual Luciferase Reporter Assay System on a Promega GloMax luminometer. All assays were performed were performed in triplicate with triplicate independent readings (n = 3).

### Transcription factor binding predictions

Transcription factor binding analysis was using QBiC-Pred (*55*) and selecting all transcription factor families and using a p-value threshold of 0.0001 and output to a VCF format. Once predictions were obtained, transcription factor were then cross referenced with RNA sequencing from mouse embryonic brains at day 11.5 to identify which transcription factors are highly expressed.

## Acknowledgments

Thank you to Dr. Evan Eichler, Dr. Jay Shendure, Dr. Barak Cohen, and Dr. Jeffrey Kidd for helpful discussions on this work.

## Author contributions

E.M.P. and T.N.T. designed the study; E.M.P. and T.N.T. wrote the paper; E.M.P., T.J.H., M.B., N.S., A.S.A., M.C.Z., and T.N.T. assessed *de novo* mutation data; T.J.H., N.S., and A.S.A. optimized the fitDNM model for noncoding regions; E.M.P. and T.N.T. assessed morbidity map data; E.M.P., R.M., G.N., A.A., J.K., J.N., M.C.Z., and T.N.T. generated the QuicK-mer2 files and performed genotyping; E.M.P. and T.N.T. performed statistical assessment of protein-coding DNVs in *EBF3*; E.M.P., Z.C., D.U.G., and T.N.T. assessed mouse ENCODE data; E.M.P. and T.N.T. assessed the ChIP sequencing data; E.M.P. and T.N.T. assessed the RNAseq data; E.M.P., S.C., L.F., and T.N.T. designed and implemented the luciferase assays; E.M.P. and T.N.T. performed transcription factor binding analysis; B.M., R.H., J.A., D.E.D., L.A.P., and T.N.T. designed and implemented the mouse mutagenesis assays; Z.C. and D.U.G. performed Hi-C analysis; and R.A.B. assessed phenotypes of individuals with hs737 variants.

## Data and materials availability

The *de novo* callset is available at SFARI Base accession: SFARI_SSC_WGS_2a, https://base.sfari.org/. We are grateful to all of the families at the participating SSC sites, as well as the principal investigators (A. Beaudet, R. Bernier, J. Constantino, E. Cook, E. Fombonne, D. Geschwind, R. Goin-Kochel, E. Hanson, D. Grice, A. Klin, D. Ledbetter, C. Lord, C. Martin, D. Martin, R. Maxim, J. Miles, O. Ousley, K. Pelphrey, B. Peterson, J. Piggot, C. Saulnier, M. State, W. Stone, J. Sutcliffe, C. Walsh, Z. Warren, and E. Wijsman). We appreciate obtaining access to phenotypic and genetic data on SFARI Base. Approved researchers can obtain the SSC population dataset described in this study (https://www.sfari.org/resource/simons-simplex-collection/) by applying at https://base.sfari.org.

## Supplemental figure legends

**Figure S1.**
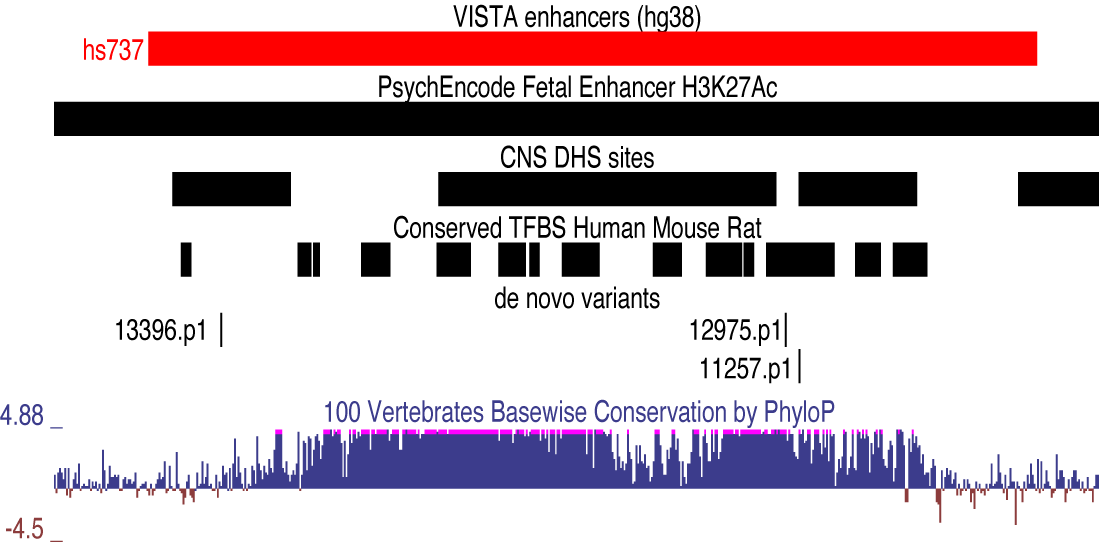
Zoom in on the hs737 enhancer with annotations from other datasets. The enhancer is in a PsychEncode fetal enhancer, contains central nervous system DNaseI hypersensitive sites, contains conserved transcription factor binding sites, and is highly conserved across the vertebrate lineage. Also shown are the locations of the hs737 *de novo* mutations identified in individuals with autism.

**Figure S2.**
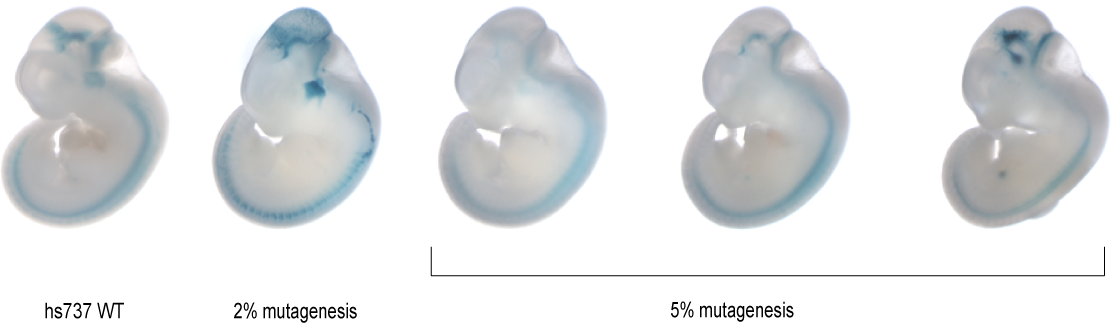
Representative mice from hs737 mutagenesis experiments.

## References

1. T. A. Manolio et al., Finding the missing heritability of complex diseases. Nature 461, 747–753 (2009).

2. K. S. Pollard, M. J. Hubisz, K. R. Rosenbloom, A. Siepel, Detection of nonneutral substitution rates on mammalian phylogenies. Genome research 20, 110–121 (2010).

3. K. J. Karczewski et al., The mutational constraint spectrum quantified from variation in 141,456 humans. Nature 581, 434–443 (2020).

4. J. E. Moore et al., Expanded encyclopaedias of DNA elements in the human and mouse genomes. Nature 583, 699–710 (2020).

5. J. Grove et al., Identification of common genetic risk variants for autism spectrum disorder. Nature genetics 51, 431–444 (2019).

6. T. N. Turner et al., Genomic Patterns of De Novo Mutation in Simplex Autism. Cell 171, 710-722.e712 (2017).

7. J. Y. An et al., Genome-wide de novo risk score implicates promoter variation in autism spectrum disorder. Science (New York, N.Y.) 362, (2018).

8. J. Zhou et al., Whole-genome deep learning analysis reveals causal role of noncoding mutations in autism. bioRxiv, (2018).

9. E. Markenscoff-Papadimitriou et al., A Chromatin Accessibility Atlas of the Developing Human Telencephalon. Cell 182, 754-769.e718 (2020).

10. I. Iossifov et al., The contribution of de novo coding mutations to autism spectrum disorder. Nature, (2014).

11. B. J. O’Roak et al., Exome sequencing in sporadic autism spectrum disorders identifies severe de novo mutations. Nature genetics 43, 585–589 (2011).

12. B. J. O’Roak et al., Sporadic autism exomes reveal a highly interconnected protein network of de novo mutations. Nature 485, 246–250 (2012).

13. S. De Rubeis et al., Synaptic, transcriptional and chromatin genes disrupted in autism. Nature 515, 209–215 (2014).

14. S. J. Sanders et al., De novo mutations revealed by whole-exome sequencing are strongly associated with autism. Nature 485, 237–241 (2012).

15. S. Sandin et al., The heritability of autism spectrum disorder. Jama 318, 1182–1184 (2017).

16. D. Levy et al., Rare de novo and transmitted copy-number variation in autistic spectrum disorders. Neuron 70, 886–897 (2011).

17. J. Sebat et al., Strong association of de novo copy number mutations with autism. Science (New York, N.Y.) 316, 445–449 (2007).

18. S. J. Sanders et al., Multiple recurrent de novo CNVs, including duplications of the 7q11.23 Williams syndrome region, are strongly associated with autism. Neuron 70, 863–885 (2011).

19. S. Girirajan et al., Refinement and discovery of new hotspots of copy-number variation associated with autism spectrum disorder. American journal of human genetics 92, 221–237 (2013).

20. I. Iossifov et al., De novo gene disruptions in children on the autistic spectrum. Neuron 74, 285–299 (2012).

21. B. M. Neale et al., Patterns and rates of exonic de novo mutations in autism spectrum disorders. Nature 485, 242–245 (2012).

22. T. N. Turner et al., Genome Sequencing of Autism-Affected Families Reveals Disruption of Putative Noncoding Regulatory DNA. American journal of human genetics 98, 58–74 (2016).

23. W. M. Brandler et al., Paternally inherited cis-regulatory structural variants are associated with autism. Science (New York, N.Y.) 360, 327–331 (2018).

24. K. A. Frazer, L. Pachter, A. Poliakov, E. M. Rubin, I. Dubchak, VISTA: computational tools for comparative genomics. Nucleic acids research 32, W273–279 (2004).

25. A. Visel, S. Minovitsky, I. Dubchak, L. A. Pennacchio, VISTA Enhancer Browser--a database of tissue-specific human enhancers. Nucleic acids research 35, D88–92 (2007).

26. Y. Jiang et al., Incorporating Functional Information in Tests of Excess De Novo Mutational Load. American journal of human genetics 97, 272–283 (2015).

27. H. Guo et al., Inherited and multiple de novo mutations in autism/developmental delay risk genes suggest a multifactorial model. Molecular autism 9, 64 (2018).

28. E. K. Ruzzo et al., Inherited and De Novo Genetic Risk for Autism Impacts Shared Networks. Cell 178, 850-866.e826 (2019).

29. A. B. Wilfert et al., Recent ultra-rare inherited mutations identify novel autism candidate risk genes. bioRxiv, 2020.2002.2010.932327 (2020).

30. I. Jung et al., A compendium of promoter-centered long-range chromatin interactions in the human genome. Nature genetics 51, 1442–1449 (2019).

31. B. W. van Bon et al., Disruptive de novo mutations of DYRK1A lead to a syndromic form of autism and ID. Molecular psychiatry, (2015).

32. J. Kaplanis et al., Integrating healthcare and research genetic data empowers the discovery of 28 novel developmental disorders. bioRxiv, 797787 (2020).

33. B. P. Coe et al., Refining analyses of copy number variation identifies specific genes associated with developmental delay. Nature genetics 46, 1063–1071 (2014).

34. G. M. Cooper et al., A copy number variation morbidity map of developmental delay. Nature genetics 43, 838–846 (2011).

35. F. Shen, J. M. Kidd, Rapid, Paralog-Sensitive CNV Analysis of 2457 Human Genomes Using QuicK-mer2. Genes (Basel) 11, (2020).

36. S. J. Sanders et al., Insights into Autism Spectrum Disorder Genomic Architecture and Biology from 71 Risk Loci. Neuron 87, 1215–1233 (2015).

37. R. L. Collins et al., A structural variation reference for medical and population genetics. Nature 581, 444–451 (2020).

38. S. K. Reilly et al., Evolutionary genomics. Evolutionary changes in promoter and enhancer activity during human corticogenesis. Science (New York, N.Y.) 347, 1155–1159 (2015).

39. W. J. Kent et al., The human genome browser at UCSC. Genome research 12, 996–1006 (2002).

40. D. U. Gorkin et al., An atlas of dynamic chromatin landscapes in mouse fetal development. Nature 583, 744–751 (2020).

41. J. Ernst, M. Kellis, ChromHMM: automating chromatin-state discovery and characterization. Nature methods 9, 215–216 (2012).

42. C. A. Sloan et al., ENCODE data at the ENCODE portal. Nucleic acids research 44, D726–732 (2016).

43. J. R. Dixon et al., Topological domains in mammalian genomes identified by analysis of chromatin interactions. Nature 485, 376–380 (2012).

44. B. Bonev et al., Multiscale 3D Genome Rewiring during Mouse Neural Development. Cell 171, 557-572.e524 (2017).

45. N. C. Durand et al., Juicer Provides a One-Click System for Analyzing Loop-Resolution Hi-C Experiments. Cell Syst 3, 95–98 (2016).

46. B. P. Coe et al., Neurodevelopmental disease genes implicated by de novo mutation and copy number variation morbidity. Nature genetics 51, 106–116 (2019).

47. J. S. Ware, K. E. Samocha, J. Homsy, M. J. Daly, Interpreting de novo variation in human disease using denovolyzeR. Current protocols in human genetics 87, 7.25.21-15 (2015).

48. H. Sleven et al., De Novo Mutations in EBF3 Cause a Neurodevelopmental Syndrome. American journal of human genetics 100, 138–150 (2017).

49. H. T. Chao et al., A Syndromic Neurodevelopmental Disorder Caused by De Novo Variants in EBF3. American journal of human genetics 100, 128–137 (2017).

50. F. L. Harms et al., Mutations in EBF3 Disturb Transcriptional Profiles and Cause Intellectual Disability, Ataxia, and Facial Dysmorphism. American journal of human genetics 100, 117–127 (2017).

51. F. Hormozdiari, O. Penn, E. Borenstein, E. E. Eichler, The discovery of integrated gene networks for autism and related disorders. Genome research 25, 142–154 (2015).

52. L. A. Pennacchio et al., In vivo enhancer analysis of human conserved non-coding sequences. Nature 444, 499–502 (2006).

53. S. Chatterjee et al., Enhancer Variants Synergistically Drive Dysfunction of a Gene Regulatory Network In Hirschsprung Disease. Cell 167, 355-368.e310 (2016).

54. J. Zhao, D. Li, J. Seo, A. S. Allen, R. Gordân, Quantifying the Impact of Non-coding Variants on Transcription Factor-DNA Binding. Res Comput Mol Biol 10229, 336–352 (2017).

55. V. Martin, J. Zhao, A. Afek, Z. Mielko, R. Gordân, QBiC-Pred: quantitative predictions of transcription factor binding changes due to sequence variants. Nucleic acids research 47, W127–w135 (2019).

56. G. D. Fischbach, C. Lord, The Simons Simplex Collection: a resource for identification of autism genetic risk factors. Neuron 68, 192–195 (2010).

57. P. B. Celestino-Soper et al., Use of array CGH to detect exonic copy number variants throughout the genome in autism families detects a novel deletion in TMLHE. Human molecular genetics 20, 4360–4370 (2011).

58. T. Gaugler et al., Most genetic risk for autism resides with common variation. Nature genetics 46, 881–885 (2014).

59. D. M. Werling et al., An analytical framework for whole-genome sequence association studies and its implications for autism spectrum disorder. Nature genetics 50, 727–736 (2018).

60. R. Bernier et al., Disruptive CHD8 mutations define a subtype of autism early in development. Cell 158, 263–276 (2014).

61. R. K. Earl et al., Clinical phenotype of ASD-associated DYRK1A haploinsufficiency. Molecular autism 8, 54 (2017).

62. The Genotype-Tissue Expression (GTEx) project. Nature genetics 45, 580–585 (2013).

63. A. R. Quinlan, I. M. Hall, BEDTools: a flexible suite of utilities for comparing genomic features. Bioinformatics (Oxford, England) 26, 841–842 (2010).

64. T. N. Turner et al., Sex-Based Analysis of De Novo Variants in Neurodevelopmental Disorders. American journal of human genetics 105, 1274–1285 (2019).

65. W. Huang da, B. T. Sherman, R. A. Lempicki, Systematic and integrative analysis of large gene lists using DAVID bioinformatics resources. Nature protocols 4, 44–57 (2009).

66. J. Ren et al., DOG 1.0: illustrator of protein domain structures. Cell Res 19, 271–273 (2009).

67. C. Tokheim et al., Exome-Scale Discovery of Hotspot Mutation Regions in Human Cancer Using 3D Protein Structure. Cancer research 76, 3719–3731 (2016).

68. N. Niknafs et al., MuPIT interactive: webserver for mapping variant positions to annotated, interactive 3D structures. Human genetics 132, 1235–1243 (2013).

